# DCLRE1B/hSNM1B (Apollo) is not acutely required for human pluripotent stem cell survival

**DOI:** 10.1101/2023.07.29.551071

**Authors:** Rebecca Bartke, Dirk Hockemeyer

## Abstract

Telomeric DNA ends in a 3’ single stranded overhang that is implicated in the protective function of telomeres ensuring genomic stability in mammals. Telomere overhang formation relies on the coordinated interplay between DNA synthesis and exonuclease activity. DCLRE1B/hSNM1B/Apollo generates an initial resection at the newly synthesized, blunt-ended leading strand telomere. This resection is thought to be required for further nucleolytic processing at the leading strand telomere. Here, we investigated the functional relevance of Apollo in human pluripotent stem cells (hPSCs) by generating Apollo deficient cells. Leveraging CRISPR/Cas9 technology, we generated locally haploid hPSCs (loHAPs) that lack one allele of Apollo. Subsequently, we mutated the remaining Apollo allele and monitored the resultant allele spectrum over 3 weeks. Surprisingly, cells survived regardless of Apollo status. These results suggest that, in hPSCs, Apollo is not acutely essential for cellular survival.

## Introduction

Telomeres, the caps at the ends of linear chromosomes, play a crucial role in maintaining genome integrity and cell viability (Reviewed in ^1^). In most human cells, the length of the telomere decreases with each cell division; stem cells and most cancer cells, however, maintain telomere length through addition of telomeric repeats by the enzyme telomerase (Reviewed in ^2–5^). This telomere attrition rate in cells without telomerase could affect cellular aging and tumor suppression ^6^. Thus, telomere length regulation is critical for the prevention of cancer and the maintenance of adult stem cell reserves (Reviewed in ^2–5^). Telomeres terminate with a 3’ overhang comprising 50-400 nucleotides of G-rich repeats ^7,8^. The size of the overhang is proposed to be correlated with the telomere shortening rate in telomerase-deficient human cells ^9^. Both the shelterin component POT1 and telomerase bind the overhang, potentially modulating the elongation and protection of telomeric DNA ^10–13^. Additionally, the overhang contributes to the formation of a protective structure known as the t-loop, which serves to suppress the activation of DNA damage repair pathways ^14,15^..

The overhang is found on both leading and lagging strand telomeres (^7,8^. DNA synthesis at the lagging strand leaves a single stranded DNA overhang; DNA synthesis at the leading strand terminates in a blunt end ^16^. Further research in mouse embryonic fibroblasts supports a model in which nucleolytic activity and DNA fill-in synthesis coordinate telomere maturation (Lam et al., 2010; Wu et al., 2010, 2012). Shelterin accessory factor hSNM1B/DCLRE1B/Apollo nuclease, referred to here as Apollo, interacts with shelterin component TRF2, which binds double stranded DNA (Bae et al., 2008; Chen et al., 2008; Lenain et al., 2006; van Overbeek & de Lange, 2006). Exonuclease1 then transiently elongates all overhangs through resection. Finally, the CTC1-STN1-TEN1 (CST) complex and DNA Pol Alpha primase work together to perform fill-in synthesis, resulting in telomeres and overhangs of appropriate length ^21–23^. In agreement, disruption of the TRF2-Apollo interaction led to increased leading-strand to leading-strand telomere fusions and DNA damage signaling at telomeres (Chen et al., 2008; Lam et al., 2010; Lenain et al., 2006; Wu et al., 2010; Ye et al., 2010). Inhibiting the NHEJ pathway with Ku70 deletion rescued this telomere dysfunction in mouse embryonic fibroblasts (Lam et al., 2010).

As a shelterin accessory factor, Apollo has roles outside of telomere maintenance (Reviewed in Schmiester & Demuth, 2017). Apollo deficient cells were more sensitive to interstrand crosslink-inducing agents MMC and cisplatin, suggesting that Apollo may have a role in interstrand crosslinking (ICL) repair (Ishiai et al., 2004; Nojima et al., 2005). Irradiation sensitivity has been observed in siRNAmediated hSNM1B/Apollo depleted HeLa cells ^25^. However, after radiation of shRNA-mediated Apollo depleted HEK293, no increased sensitivity to radiation was observed ^17^. Additionally, Apollo may have a role in repairing stalled replication forks: Apollo was dispensable for cell recognition of stalled replication forks, but catalytically active Apollo was required for recruiting FANCD2 and BRCA1, which protect the stalled replication forks from degradation during repair (Mason et al., 2013). Furthermore, Apollo may also have a role in relieving topological stress as overexpression of Apollo rescued the deleterious effects of topoisomerase 2-alpha depletion (Ye et al., 2010). Expression of a human patient variant unable to bind TRF2 in immortalized fibroblast cells did not result in increased sensitivity to ICL inducing agents, but did exhibit telomere dysfunction ^26^.

Because a functional overhang is thought to be required for cell survival, and Apollo is thought to be required for a functional overhang, it is interesting that it is possible to generate Apollo deficient mouse embryos ^15,27,28^. However, the embryos die before maturation into adulthood, indicating Apollo is essential for development ^27,27^. In humans, inherited Apollo deficiency causes severe bone marrow failure and developmental defects ^26,29^. Another 2 patient variants of Apollo had mutations in the catalytic domain of Apollo; cells from these patients exhibited telomere fusions, but normal global telomere length ^29^. In contrast, HT-1080 Apollo knockout cells exhibited excessive telomere shortening and telomere fusions ^29^. To explain the differences in phenotypes among murine, HT-1080 cancer cells, patients with mutations in Apollo, the authors propose that the patient mutations are hypomorphic alleles ^29^. Indeed, the phenotypes of Apollo depletion also vary among cell lines ^17,25^.

Here, we attempted functional dissection of Apollo through genetic dissection in human pluripotent stem cells.

## Results

### Locally haploid human pluripotent stem cell line (LoHAP) generation to interrogate Apollo function

We recently developed an experimental system to efficiently evaluate mutations in human pluripotent stem cells (hPSCs). This approach excises one copy of a gene by paired CRISPR cutting. This results in a haploid setting at this region while all other loci within the genome remain diploid. We refer to these cells as locally haploid human pluripotent stem cells (LoHAPs) ^30^. In loHAPs, mutations of interest can be introduced into the remaining allele, all under endogenous control. From this, a strict genotype to phenotype relationship can be determined in a genetically stable cell system, without potentially obfuscating heterozygous or compound heterozygous mutations.

Apollo LoHAP cell lines were made twice independently, first in a CDKN2A null background and then in a wildtype background (editing efficiencies between <1% and 4.6%) ^31,32^. Successful generation of the LoHAPs was confirmed using a junction PCR as well as a PCR spanning a SNP present in WIBR#3 at the Apollo locus (Figure 1). We successfully generated both allelic variants based on this PCR. Survival in culture for at least 10 passages demonstrated the haplosufficiency of Apollo.

**Figure 1.**
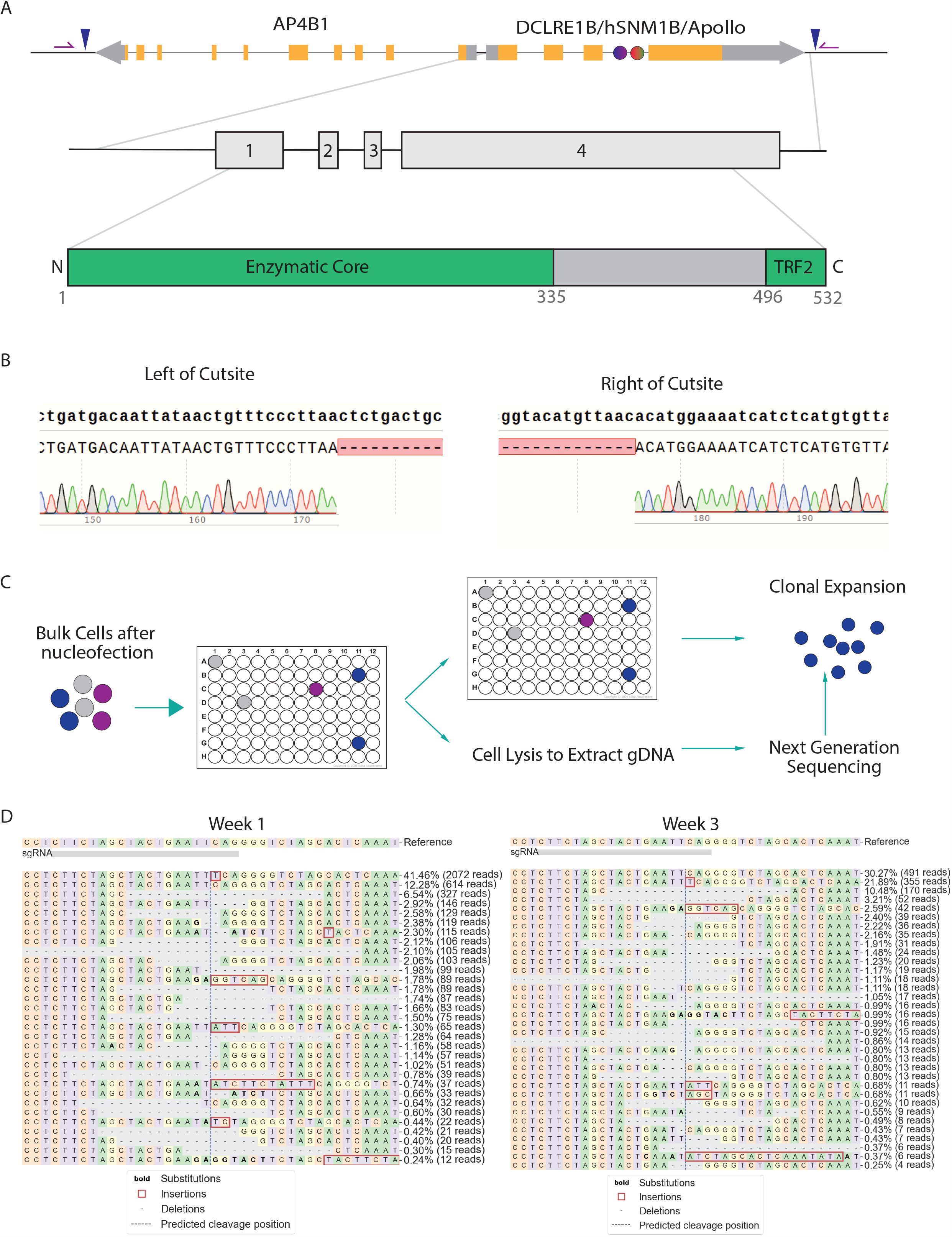
Generation of the Apollo LoHAP. (A) Schematic of Apollo transcript. Blue triangles represent flanking CRISPR/Cas9 for gene excision. Purple arrows represent primers for genotyping. Circles represent SNP locations used for further genotyping. (B) After editing, PCR is performed. As the region spanning the two primers in unedited cells is over 20kb, the only PCR products generated are from cells with successful editing events, where the resulting PCR band is likely <300 bp. Exact band sizes could change due to imprecise lesion repair. (C) Limited dilution as described in methods below. (D) CRISPResso2 alignment and visualization of a biological replicate at the first and last passage after CRISPR/Cas9 editing with an sgRNA targeting upstream of TRF2 interacting domain ^43^.

### Apollo knockout human pluripotent stem cells persist in culture

To understand which of Apollo’s functional domains are required, if any, we mutated the remaining Apollo allele in Apollo LoHAPs with CRISPR/Cas9 (Figure 2). When the CRISPR/Cas9 cut, resulting NHEJ can result in an allele spectrum caused by indels. If a particular amino acid is required for cell viability, and it is excised, we expect to see its allele frequency to deplete. Similarly, if a domain is essential, we expect that persisting alleles must have that domain intact. We previously utilized this approach for BRCA2, demonstrating that functional alleles are maintained over 3 weeks, while mutations which negatively affect cellular viability deplete in allele frequency over time ^30^.

**Figure 2.**
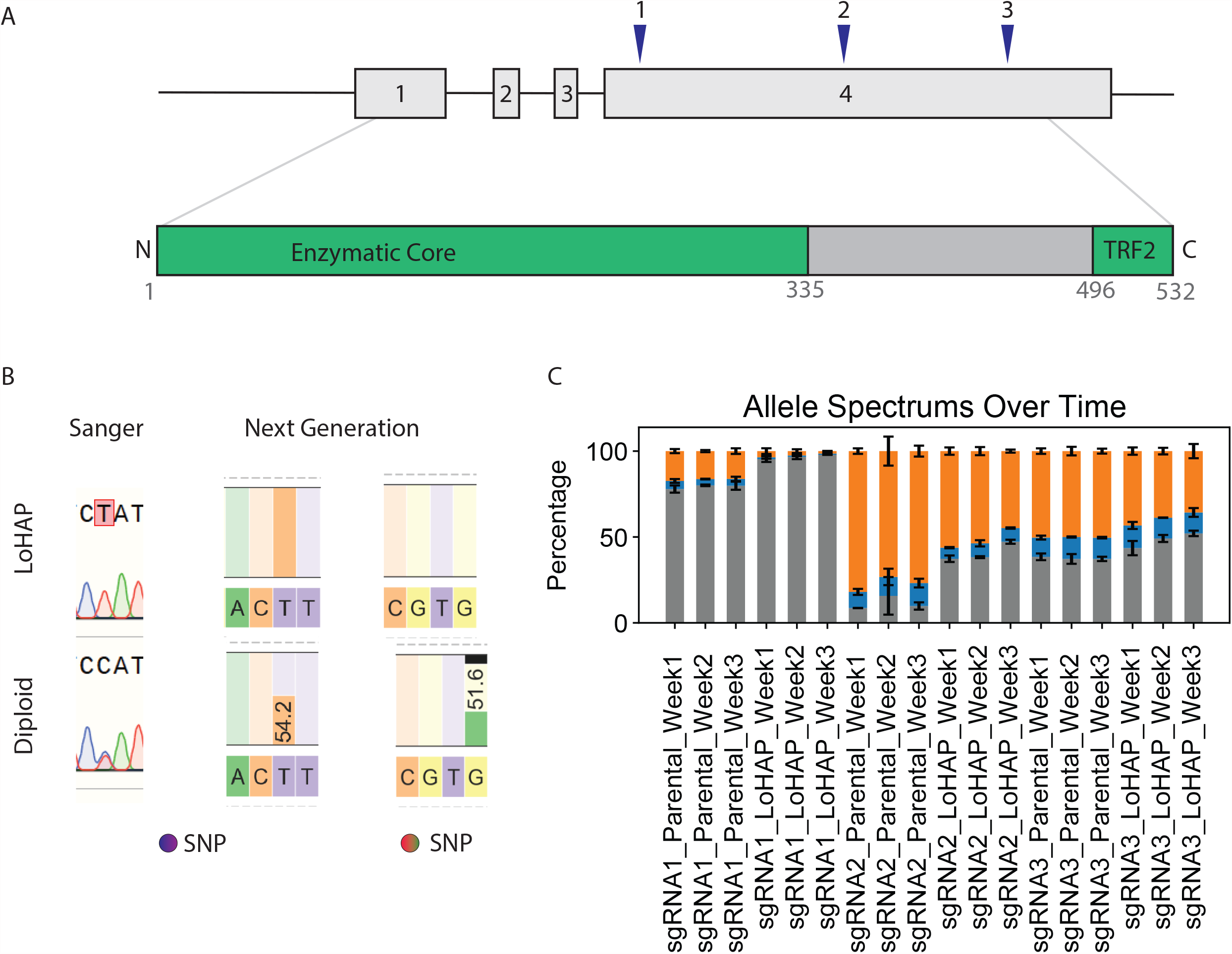
Targeting the remaining Apollo allele. (A) Schematic of guide positions at the endogenous locus. (B) SNP genotyping PCR performed over the remaining allele. (C) Relative percentages of out-of-frame and in-frame and unedited alleles after targeting over time, as determined by NGS. Gray, blue, orange, correspond to unedited, in-frame, and out-of-frame, respectively. Each value corresponds to 3 independent biological replicates. Error bars are standard deviation.

Guides to mutate Apollo were designed to cut within Exon 4, the final and largest exon of Apollo ^25,33^. This exon is also the only exon shared by all transcript variants ^25,33^. Mutations in Apollo F504, L506, and P508 abolish its interaction with shelterin component TRF2, and a crystal structure of TRF2_TRFH_-Apollo_496-532_ identifies Apollo residues 498-509 at the TRF2-Apollo interaction surface ^18^. Mutations in the HxHxDH motif and H230 disrupt Apollo catalytic function ^34^. We designed guides to dissect each of Apollo’s proposed functions: 1) catalytic function, 2) combined function, and 3) telomere-specific function. The first guide RNA targeted the beginning of exon 4, by S183. The second guide RNA targeted the center of Exon 4, near K371. The third guide, positioned at the closest available NGG PAM to the TRF2 binding site, targeted P508. Through a comparison of the allele spectra created by these guides, we aimed to decipher which of Apollo’s functions are required for continued cell proliferation.

These guides were individually nucleofected into their respective LoHAPs and diploid parentals. Next, the allele spectra of 3 independent replicates were tracked over three weeks (Figure 1D). Sequencing, filtering, and alignment of these spectra revealed indel scarring of assorted sizes around the cutsites. Deletions of 3, 6, or 9 nucleotides were presumed to lead to in-frame mutations, such that only local amino acid composition was changed. Deletion sizes indivisible by 3 were presumed to have led to frameshift mutations. Alleles with these deletion sizes were presumed to either lead to unstable transcripts or truncated proteins. Through a comparison of which categories of alleles were tolerated at each site, we aimed to understand which of Apollo’s functions were critical for cell survival.

Surprisingly, both in-frame and out-of-frame mutations were created at all sites and persisted over time in the parental and the Apollo LoHAP, regardless of position (Figure 2). This suggests that neither Apollo’s enzymatic core functions nor its telomere-specific functions are acutely required in human pluripotent stem cells.

## Discussion

The persistence of out-of-frame mutations persisting in Apollo was initially unexpected, as loss of function mutations for Apollo are incompatible with murine life, and there are no known patients lacking all Apollo expression ^27,28^. Cells with telomere fusions or as few as five deprotected telomeres exit the cell cycle through activation of ATM/ATR DNA damage repair machinery ^35^. Fibroblast cells with these Apollo variants showed increased sensitivity to DNA damage causing agents, increased frequency of extremely short telomeres, and G-overhang formation defects ^29^. HT-1080 Apollo knockout cells exhibited excessive telomere shortening and telomere fusions ^29^. HT-1080 Apollo knockout cells did persist in cell culture ^29^. Similarly, here, hPSC Apollo knockout cells persist.

Recent investigations have revealed the dispensability of TRF2, the mediator between Apollo and the telomere, in mouse embryonic stem cells which exhibit transcriptional similarities to the 2-cell state (Markiewicz-Potoczny et al., 2021). These cells did exhibit DNA damage signaling, but no fusions (Markiewicz-Potoczny et al., 2021). Relevantly, out-of-frame mutations resulting in a loss of the TRF2 binding site in Apollo were also tolerated. One interpretation of this is that Apollo loss in hPSCs may result in DNA damage signaling, but this DNA damage signaling does not lead to exiting the cell cycle. LoHAPs can be differentiated into cell types of interest ^30^. Future studies may differentiate Apollo knockout LoHAPs, to understand the contribution of cell type to Apollo phenotype.

Another possibility is that Apollo could instead serve an essential role in telomere length maintenance. In this scenario, Apollo deficient cells could be experiencing gradual telomere shortening. In our experiment we only sampled cells at a maximum of 3 weeks after editing. hPSCs have a telomeric reserve, and can take 8-10 weeks until telomere shortening defects result in cell death ^36^. This hypothesis could explain why generating Apollo knockout mouse embryos is possible, but Apollo knockout adult mice ^27,28^. Future experiments would extend the sampling window of this experiment and examine telomere length changes over time, and in differentiated cells, to determine when Apollo is required for telomere length maintenance.

## Methods and Materials

### Cell culture maintenance

Cell engineering was performed in WIBR3 hESCs, NIH stem cell registry #0079. hESCs were maintained on a monolayer of CD-1 strain mouse embryonic fibroblasts (MEFs). hESC maintenance media was made up of DMEM/F12 [Gibco], supplemented with 20% KnockOut Serum Replacement [Gibco], 1mM glutamine [Sigma Aldrich], 1% non-essential amino acids [Gibco], 100U/mL Penicillin-Streptomycin, 0.1 mM β-mercaptoethanol [Sigma-Aldrich], and 4ng/mL heat-stable FGF-Basic (AA 1-155) [Gibco]. hESC wash media was DMEM/F12 [Gibco], 5% heat inactivated Newborn Calf Serum [5% Sigma-Aldrich], 100U/mL Penicillin-Streptomycin. hESC maintenance media was replaced daily. On most weeks, cells were passaged every 5-7 days enzymatically with 1.5mg/mL collagenase type IV [Gibco]. Cells were collected, washed, and sedimented at least 3 times in hESC wash media, then replated in hESC maintenance media at a ratio of 1:6. However, on weeks where maintaining allele complexity was paramount like in the allele tracking experiments above, cells were passaged as single cells using 0.25% Trypsin-EDTA. The day before, the day of, and the day after passaging using trypsin, hESC maintenance media was supplemented with Rock Inhibitor Y-27632.

### Generation of LoHAPs

The following is summarized our paper ^30^, based on previously reported gene editing strategies for large scale deletions ^37–40^. To excise a copy of a gene of interest using the LoHAP method, sgRNA were designed to flank the gene of interest. sgRNA were designed from the end of the previous gene to the end of the gene-of-interest, to limit unintended transcriptional regulation changes on proximal genes ^30^. Guide RNAs with high predicted specificity scores were chosen ^41,42^. After nucleofection, cells were seeded onto 96 well plates at either 1,000 cells/well, or 100 cells/well. Higher density seeding permits capture of more alleles per well; when trying to capture a rare editing event, seeding first at higher density, then performing limited dilution on the enriched well can ease clone isolation. Remaining cells were replated onto MEFs, and DNA extracted after 3 days to act as positive control for the junction PCR (Figure 1B, 1C).

Media on the 96 well plates was changed at days 4, 7, 10, and 12. The day before, the day of, and the day after passaging using trypsin, hESC maintenance media was supplemented with Rock Inhibitor Y-27632. Cells were then replica plated as follows: 100uL PBS-wash, 40uL trypsin for 5 minutes at 37 °C, addition of 60uL FBS/hPSC media (replacing 10% KSR with 10% FBS in regular hPSC media), trituration. Then, this 100 uL was split into two 96-well plates. The first plate had MEFs and continued to be cultured, with media changes as above. The second plate had 50 μl 2x lysis buffer (100mM KCl, 4mM MgCl2, 0.9% IGEPAL, 0.9% Tween-20, 500 μg/ml proteinase K, in 20mM Tris pH 8) for DNA extraction. The cell lysate was incubated overnight at 50°C, then heated for 10 minutes at 95°C.

Junction PCR was performed using primers flanking the guide RNAs’ target sites, using either Titan or PrimeStarGXL polymerases, and 2uL of cell lysate (Figure 1B, 1C). Amplicons were resolved with 1% agarose gel electrophoresis, followed by sequencing by Next Generation Sequencing (NGS) or Sanger sequencing. To confirm heterozygosity, two single-nucleotide polymorphisms (SNP) were identified within the deletion region from whole genome sequencing data (Figure 1A). Primers containing NGS barcode (5’ GCTCTTCCGATCT 3’) flanked SNP positions in clones with identified junction PCR bands. Amplicons were purified using SPRI bead purification at the UC Berkeley DNA Sequencing Facility, i5/i7 barcoded, pooled, and run on 150 PE iSEQ. NGS was analyzed using CRISPResso2 to perform computational quality control and alignment ^43^. Wells of interest with a single nucleotide remaining at a SNP were then further subcloned by low-density seeding or manual picking to ensure clonality (Figure 2B).

### Cas9-RNP nucleofection for Essentiality Test

hPSCs were cultured on MEF prior to nucleofection. They were separated from the MEF layer with 1mg/mL collagenase for 20 minutes. Colonies were suspended and the collagenase was neutralized by collecting the suspension with 10 x volume of Wash Media (DMEM/F12, 5% FBS, 1X Penicillin/streptomycin). After a 5 minute incubation at room temperature, the MEF and wash media suspension was aspirated, leaving hPSC colonies. 0.25% Trypsin-EDTA was added to this and incubated for 5 minutes at 37°C, then titurated with a 10X volume of Wash media. This was spun down at 1000 RPM for 5 minutes. They were washed with phosphate-buffered saline (PBS-) and spun down again at 1000 RPM for 5 minutes. While this was happening, the Cas9-RNP complex was made: 300 pMol of sgRNA (Synthego) and 80 pMol of Cas9-protein (SB3 MacroLab, UC Berkeley) were combined with nuclease-free water to a volume of 10uL, and incubated at room temperature for a minimum of 10 minutes. Half a million cells were resuspended in 20uL Lonza P3 primary nucleofection reagent, then added to the Cas9-RNP. This was then nucleofected using program CA137 on the Lonza nucleofector 4D. Finally, the nucleofected cells were resuspended into 9 mL, and replated into 3 now independent biological replicates, 3mL into each well of a 6 well plate. At 8 days, 13 days, and 18 days, 5 % of the cells were passaged onto new wells. The remaining 95% was divided into 2 tubes for freezing of cell stocks, and two tubes for DNA extraction. After DNA extraction, PCR was performed with primers flanking sgRNA cutsite to generate a 150-250 nucleotide amplicon. Amplicons were purified using SPRI bead purification at the UC Berkeley DNA Sequencing Facility, i5/i7 barcoded, pooled, and run on 150 PE iSEQ. NGS result was analyzed using CRISPResso2 ^43^.

### In-Frame/out-of-Frame experiment and analysis

A custom python script performed downstream analysis. The beginning and ends of the reads exhibit increased rates of incorrect PCR calls; to remedy this, the first and last 20 nucleotides of the reads were trimmed. To reduce the impact of PCR error on interpretation of these alleles, only DNA changes within 2 nucleotides of either a cutsite, or a lesion extending from the cutsite, were considered biological-relevant mutations. Other DNA changes, here assumed to be PCR error, were converted to wildtype sequence. Next, the size of the deletions and insertions were counted. The read counts of identical alleles were now summed together. To further reduce PCR error noise, all alleles with fewer than 5 reads were filtered out. The number of insertions and deletions were counted within each allele. An allele was considered “in-frame” if the absolute value of the difference between the number of insertions and the number of deletions was a multiple of 3, and the sequence was not wildtype. If there were no substitutions, deletions, or insertions, the allele was declared “wildtype”. Otherwise, it was considered an “out-of-frame” allele. The averages from each condition each week among the biological replicates determined the height of the bars; error bars depict the standard deviation calculated among the three biological replicates at each time point.

### Mycoplasma testing

All cells were tested at least monthly for *Mycoplasma* contamination. For *Mycoplasma* detection, genomic DNA extraction was performed with TENS Buffer (10 mM Tris HCL, pH 8, 1 mM EDTA, pH 8, 100mM NaCl; 1:40 Proteinase K was added day of use) overnight at 37°C. Then, gDNA was phenol chloroform extracted, isopropanol precipitated, and resuspended in TE Buffer. Finally, the sample was diluted 1:10 before submission.

## Author Contributions

R.B. and D.H. conceptualized the project, designed all experiments and wrote the manuscript.

R.B. performed the experiments, wrote the scripts, and analyzed the data.

## Data Availability

The raw NGS data and whole genome sequencing (WGS) data of WIBR3 are available upon request. The Python pipeline is also available upon request.

## Acknowledgments

The UC Berkeley MCB Graduate Affairs Committee requires Ph.D. students to publish a first author paper to graduate. The purpose of submitting this work to BioRxiv is to fulfill this requirement for R.B.’s graduation from UC Berkeley with her Ph.D.

## Figure Text

**Tables.**
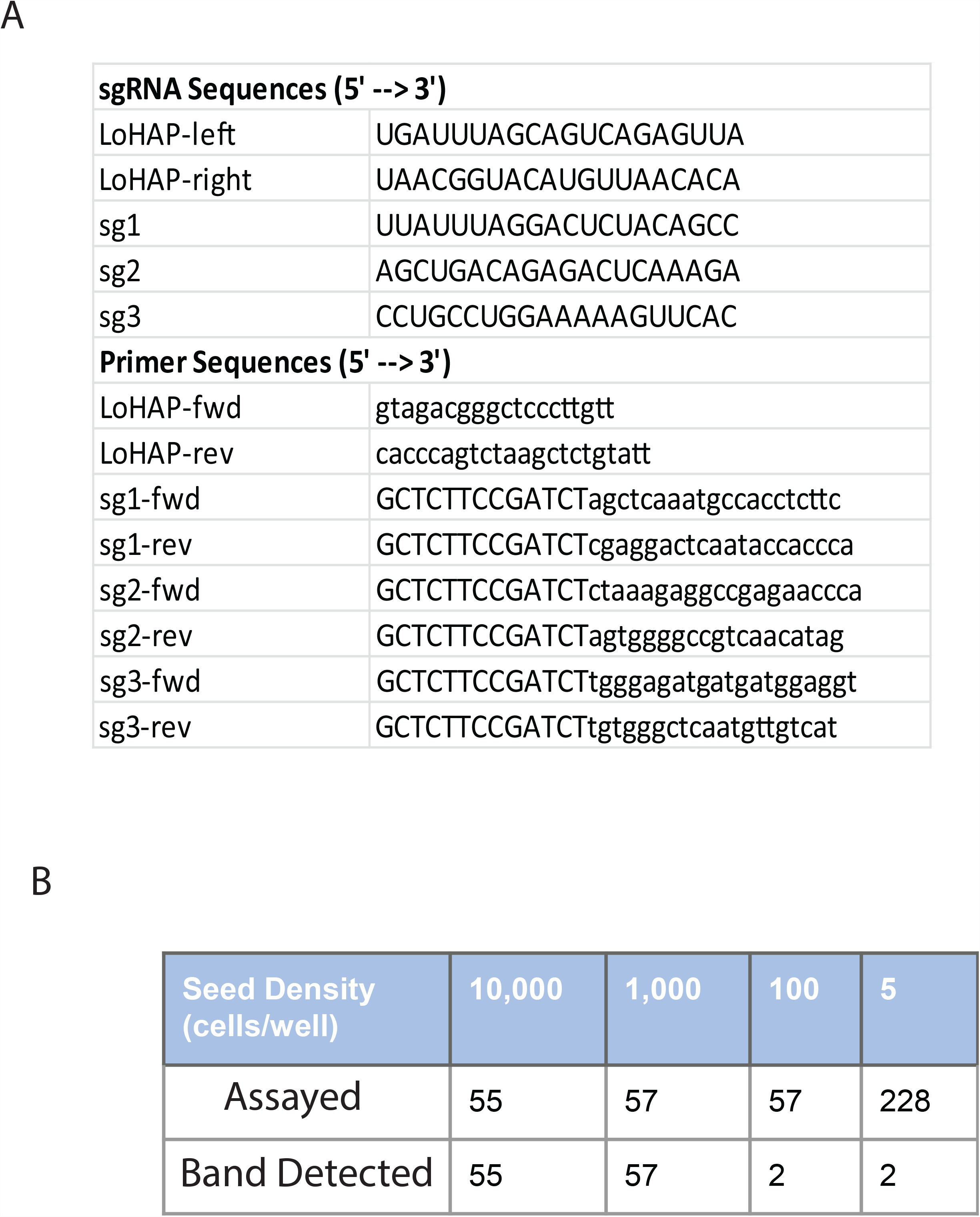
(A) sgRNA sequences used to generate the Apollo LoHAP, and primer sequences used for genotyping.(B) Representative editing efficiency for generating Apollo LoHAP in checkpoint proficient cells during limited dilution.

